# Global mRNA 3′UTR lengthening in small-cell neuroendocrine carcinoma

**DOI:** 10.64898/2026.01.26.701900

**Authors:** Yi Zhang, Xiaofan Zhao, Huan Wang, Ya-Mei Hu, Xiao-Xin Sun, Faming Zhao, Shunyu Du, Roselyn S. Dai, Nicole K. Andeen, Rosalie C. Sears, Eva Corey, Jonathan R. Brody, Joshi J. Alumkal, Gordon B. Mills, Peter S. Nelson, Mu-Shui Dai, Zheng Xia

## Abstract

Small-cell neuroendocrine carcinoma (SCNC) is a rare but highly malignant tumor subtype that primarily arises in the lung, also rarely in other organs, and as a consequence of treatment induced lineage transdifferentiation of prostate adenocarcinomas. The molecular convergence of SCNC across diverse tissues enables its identification through conserved SCNC-specific molecular markers, facilitating tumor subtype classification. As a critical post-transcriptional regulatory mechanism, alternative polyadenylation (APA) modulates 3′UTR length and significantly impacts tumor progression. However, its role in SCNC remains largely unclear. Here, we report a global 3′UTR lengthening pattern driven by APA in SCNC. We identified a set of conserved 3′UTR lengthening events across SCNCs of different tissue origins, which are strongly associated with neural development and related signaling pathways. Furthermore, we developed a neural network-based prediction model to classify SCNC by leveraging these specific APA signatures. Our study provides new insights into the post-transcriptional landscape of SCNCs and highlights APA signatures as promising biomarkers for SCNC identification.

## Introduction

Small-cell neuroendocrine carcinoma (SCNC) is an uncommon but highly malignant tumor subtype that arises in various human tissues, predominantly in the lung and also as a consequence of therapeutics that repress androgen receptor signaling in prostate cancer. SCNC is characterized by its aggressiveness and poor clinical outcomes compared to non-SCNC tumors^1–3^. For instance, lung cancer accounts for approximately 227,875 cases in the United States, with small-cell lung cancer (SCLC) constituting roughly 14% of these cases^4^. The prognosis for SCLC is poor, with a 2 year survival rate of ∼7%^2^. Current diagnostic methods for neuroendocrine tumors, including small-cell variants of the lung and prostate, primarily rely on histology, imaging, and protein biomarkers such as chromogranin A (CHGA), neural cell adhesion molecule 1 (NCAM1), and synaptophysin (SYP)^5^. However, these markers often suffer from limited sensitivity and specificity, failing to completely address diagnostic and prognostic requirements^5^. Moreover, studies indicate a molecular convergence among small-cell neuroendocrine carcinomas (SCNCs) from different tissues, where tumors increasingly resemble each other morphologically, deviating from their tissue of origin^6, 7^. Therefore, identifying conserved markers across SCNCs may be useful for early diagnosis, a refinement of tumor subclassification and the development of therapeutics.

Polyadenylation, a crucial step in messenger RNA (mRNA) maturation, adds a poly(A) tail to the 3′ untranslated region of (3′UTR) of mRNA^8^, thereby enhancing mRNA stability and translation efficiency^9, 10^. While the canonical signal AAUAAA triggers polyadenylation^11^, most genes harbor multiple polyadenylation sites, leading to alternative polyadenylation (APA)^12^. APA modulates gene expression and function by altering the inclusion of RNA- and protein-binding sites within the 3′UTR. This makes APA a potential molecular marker in various biological contexts, such as cellular senescence and tumor progression^13–16^.

While global 3’UTR shortening is a well-established hallmark of cancer relative to healthy tissues^17–20^, it remains unclear whether SCNC exhibits further distinctive APA patterns that differentiate it from other tumor subtypes. To bridge this gap, we systematically characterized the APA profiles in SCNC and non-SCNC samples from lung and prostate cancers. Strikingly, we identified a distinct global 3′UTR lengthening pattern in both small-cell lung cancer (SCLC) and neuroendocrine prostate cancer (NEPC). Leveraging this finding, we developed an APA-based prediction model utilizing a neural network framework to identify SCNC samples across cancer types. The prediction results were validated against the expression of classical SCNC marker genes and confirmed by pathological reports. Collectively, our study identifies a unique post-transcriptional regulatory mechanism in SCNC and proposes APA events resulting in lengthened transcripts as robust signatures for SCNC classification. These findings provide insights into the development and maintenance of SCNC and underscore the potential of APA as a clinical biomarker for diagnosis and prognosis.

## Results

### Global 3′UTR lengthening in SCNCs of the lung and prostate at bulk RNA-Seq level

To comprehensively characterize the APA landscape in SCNC, we employed the DaPars algorithm^18^ to quantify dynamic 3′UTR usage at the bulk RNA-Seq level. We analyzed 49 prostate cancer samples, including 34 prostate adenocarcinomas (PRAD) and 15 neuroendocrine carcinomas (NEPC) from the Beltran dataset^21^ and 192 lung cancer cell line samples, including 142 non-small cell lung cancers (NSCLC) and 50 small cell lung cancers (SCLC) from the Cancer Cell Line Encyclopedia (CCLE) dataset^22, 23^. We uncovered a global 3’UTR lengthening pattern in SCNC samples from both lung and prostate tissues and identified a set of highly conserved lengthening APA signatures to classify the SCNC samples across tissue types.

Subsequently, we developed a neural network-based prediction model using these transcript lengthening APA signatures as input to identify potential SCNC samples, which demonstrated a significant outcome difference between SCNC and non-SCNC samples (**Fig. 1A**). Specifically, based on the Percentage of Distal poly(A) site Usage Index (PDUI), we identified 451 lengthening and only 13 shortening APA events in SCLC samples compared to NSCLC samples (**Fig. 1B**). Consistently, 220 lengthening and 15 shortening APA events were identified in NEPC samples compared to PRAD samples (**Fig. 1C**). For validation, we applied APA profiling to four patient-derived xenografts (PDXs) prostate cancer samples (2 NEPCs and 2 PRADs)^24^. We observed a similar lengthening pattern in NEPC samples, further confirming the global 3’UTR lengthening trend in SCNC (**Supp. Fig 1**).

**Figure 1.**
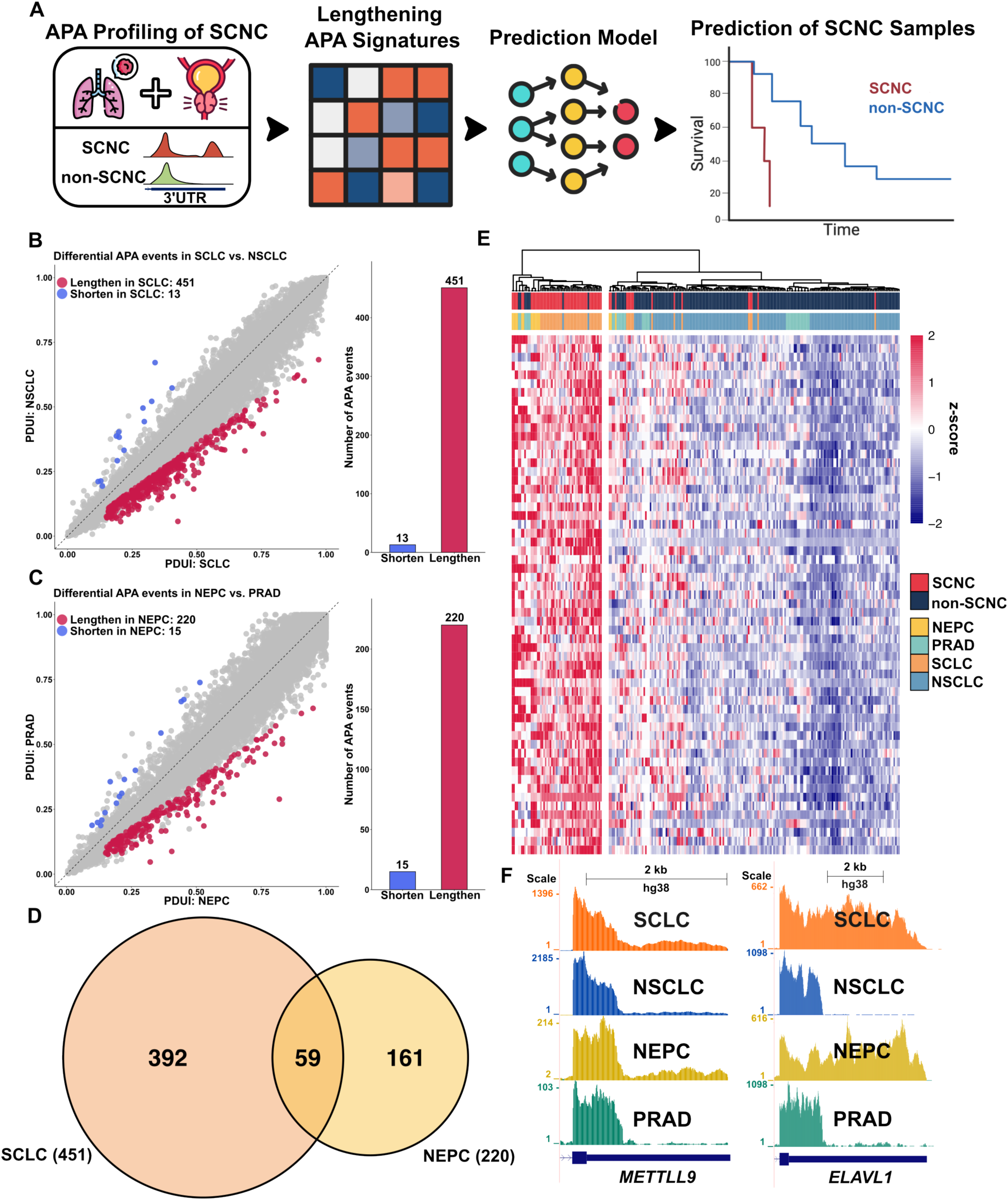
Global 3’UTR lengthening in SCNC at bulk level. **(A)** The workflow of characterizing APA landscape in SCNC samples. Targeting on lung and prostate cancer, the Percentage of Distal PAS Usage Index (PDUI) of each APA events between SCNC and non-SCNC samples were estimated to profile the APA events. A set of lengthening APA signatures was identified and the PDUI matrix of the APA signatures was utilized as the input of the Neural-Network-based prediction model. Based on the prediction model, the identified SCNC samples showed significant outcome differences comparing with non-SCNC samples. **(B)** Global 3’UTR lengthening in small cell lung cancer (SCLC). 451 lengthening and 13 shortening APA events were identified in the SCLC samples under cutoffs: FDR<0.1, |FoldChange|>=1.4, diff_PDUI>=0.05 and min_PDUI>=0.15. **(C)** Global 3’UTR lengthening in neuroendocrine prostate carcinoma (NEPC). 220 lengthening and 15 shortening APA events were identified in the NEPC samples under cutoffs: FDR<0.1, |FoldChange|>=1.4, diff_PDUI>=0.05 and min_PDUI>=0.15. **(D)** Overlapping of the lengthening APA events between SCLC and NEPC samples. 59 APA events showed a lengthening pattern between SCLC and NEPC. **(E)** Heatmap of 241 lung and prostate cancer samples clustering based on the 59 shared lengthening APA events. Each row represents one of the 59 APA events, and each column represents an individual sample. The heatmap displays z-scores calculated from the PDUI matrix, where red indicates higher usage of distal poly(A) sites (lengthening) and blue indicates lower usage of distal poly(A) sites. **(F)** Coverage plots of two representative lengthening APA events. Both METTL9 and ELAVL1 showed a longer mRNA isoforms preference in the SCNC samples from lung and prostate cancers.

Given the morphological similarity and molecular convergence of SCNC samples across tissues^5, 6^, we hypothesized that SCNC samples might share a set of lengthening APA events between lung cancer and prostate cancer, and possibly in other tissues. By intersecting the identified lengthening APA profiles, we obtained 59 shared lengthening APA events between SCLC and NEPC (**Fig. 1D**). We then used these 59 APA events to perform clustering on the 241 lung and prostate cancer samples to evaluate their capability in classifying SCNC and non-SCNC samples. Based on the PDUI matrix, we calculated the z-scores, and the resulting heatmap illustrated a distinct separation between SCNC and non-SCNC groups. Specifically, SCLC and NEPC samples exhibited consistently elevated z-scores for these shared lengthening events compared to NSCLC and PRAD samples, reflecting a conserved trend across SCNCs from lung and prostate tissues (**Fig. 1E**). A subset of SCNC samples clustered more closely with non-SCNC samples, suggesting a potential need for more precise classification via APA profiling. Finally, we selected two representative genes with lengthening APA events in both SCLC and NEPC, *METTL9* and *ELAVL1*, to demonstrate dynamic poly(A) site (PAS) usage (**Fig. 1F**). The 3’UTR lengthened transcripts of both genes were predominantly observed in the SCNC samples of lung and prostate cancers, respectively.

### Construction of the APA signature-based prediction model

The 59 APA events shared in SCLC and NEPC exhibited distinct PDUI levels between SCNC and non-SCNC samples from the lung and prostate, suggesting their potential as robust molecular markers for SCNC identification (**Fig. 1D**). To further explore the APA patterns across SCNCs from diverse tissues, we first performed a conservation analysis on these 59 events, excluding tissue-specific ones to identify universally conserved signatures across SCNC subtypes. Specifically, we evaluated their expression across 7,659 tumor samples from 25 cancer types in The Cancer Genome Atlas (TCGA). Since missing PDUI values typically indicate low gene expression or insufficient read coverage, we selected events with a missing value ratio of less than 0.01 to ensure that the identified signatures were universally measurable across diverse tumor types. In total, 20 conserved APA signatures met the criteria and were selected to train and validate the neural network model (**Fig. 2A, Supp. Fig 2**).

**Figure 2.**
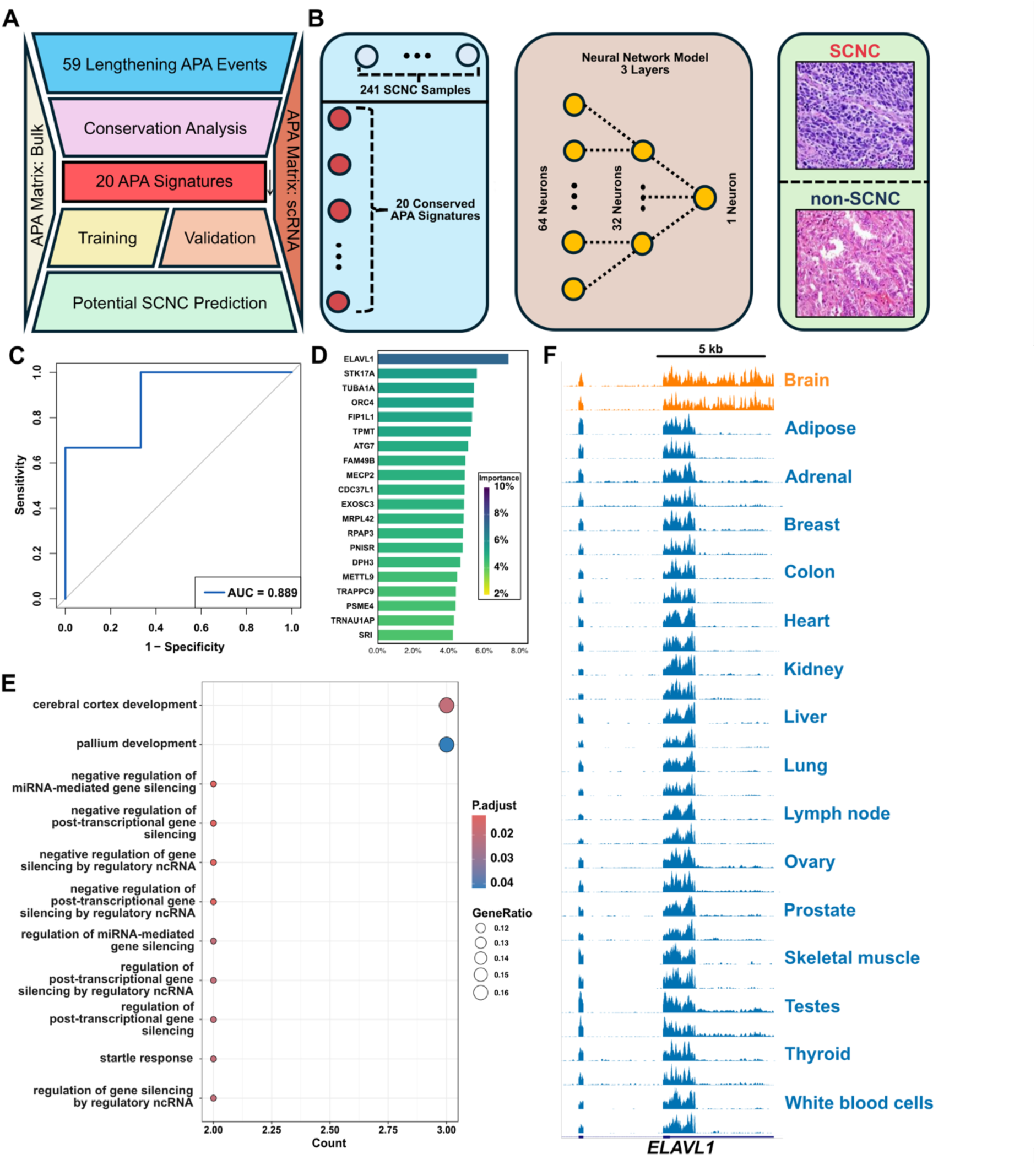
A set of APA signatures guides the prediction model based on neural network. **(A)** The workflow of establishing the prediction model. The overlapped 59 lengthening APA events between SCLC and NEPC were applied to perform the conservation analysis on 7,659 TCGA pan-cancer samples. 20 highly conserved APA signatures were identified based on the missing value frequency of each event across tissues. The matrices of the 20 APA signatures from bulk RNA-Seq and scRNA-Seq datasets were applied to perform training and validation of the prediction model. Specifically, the matrix of 28 scRNA-Seq lung cancer samples was utilized to optimize and select the cutoff for classification. The matrix of 6 scRNA-Seq prostate cancer samples was utilized to validate the prediction results. Further, the prediction model was applied to classify and predict potential SCNC samples from public datasets. **(B)** The framework of the neural-network-based framework. The input layer comprises 64 neurons, processing a 20×241 PDUI matrix derived from bulk RNA-Seq data. A hidden layer of 32 neurons performs feature reduction, followed by a single-neuron output layer for classification and prediction. Two pathology images of small cell lung cancer and non-small cell lung cancer samples serve as the examples of classification results. **(C)** ROC analysis of the prediction results of 6 scRNA-Seq samples of prostate cancer. With AUC equals 0.889, 1 NEPC (Patient #P2) and 5 non_NEPCs were identified. **(D)** Feature importance of 20 APA signatures in the prediction model, quantified using Garson’s algorithm. ELAVL1 emerged as the most influential feature in the model. **(E)** Gene Ontology analysis of the 20 APA signatures revealed enrichment in neuronal development and brain development pathways, as well as post-transcriptional regulatory processes. **(F)** Coverage plot of the 3’UTR of *ELAVL1* across 16 normal human tissues. Lengthened isoforms of *ELAVL1* were enriched in brain, indicating a correlation of APA lengthening and neural-related pathways.

The model architecture comprised an input layer with 64 neurons receiving the PDUI matrix of the 20 signatures from 241 samples (the samples used for bulk APA analysis), a hidden layer with 32 neurons for feature reduction, and an output layer with 1 neuron for binary classification. This framework generated prediction scores to determine the probability of a target sample being SCNC (**Fig. 2B**). To determine an optimal classification cutoff, we utilized pseudo-bulk PDUI matrices derived from single-cell lung cancer samples from the HTAN (Human Tumor Atlas Network) database^25^. Quality control based on epithelial tumor cell abundance yielded a high-quality cohort of 28 samples (22 SCNCs and 6 non-SCNCs) from an initial set of 52 (**Supp. Fig 3**). Using their histological tumor subtype annotations as ground truth, we determined the optimal cutoff that properly separated SCNC from non-SCNC samples. Samples with prediction scores above the cutoff were classified as SCNC, while those below were classified as non-SCNC.

For independent cross-validation, we applied the model to an independent public scRNA-Seq dataset of 6 single-cell prostate cancer samples^26^. Using the fixed cutoff derived from the lung cohort, only Patient #2 was explicitly classified as SCNC. Nevertheless, the model exhibited robust discriminative power, achieving an area under the ROC curve (AUC) of 0.889 (**Fig. 2C**). Although Patients #5 and #6 failed to cross the stringent cutoff, their prediction scores were consistently higher than Patients #1 and #4. This suggests that while factors such as single-cell data sparsity and biological heterogeneity may dampen absolute prediction scores and limit sensitivity at the strict cutoff, the model still showed correct relative ranking. Furthermore, to interpret the model’s decision-making process, we quantified feature importance using Garson’s algorithm^27^. Interestingly, *ELAVL1* emerged as the most influential feature (**Fig. 2D**), highlighting its potential driver role in SCNC biology.

Moreover, Gene Ontology (GO) analysis revealed that the 20 APA signatures were enriched in pathways related to brain development and post-transcriptional mechanisms (**Fig. 2E**). This validates the association between lengthening APA signatures and neural-related pathways, as well as the neuroendocrine biological processes. Previous studies have demonstrated that the 3′UTR lengthening is associated with neurogenesis^28, 29^, supporting the neural relevance of the APA signatures we identified. To further validate the 3′UTR lengthening pattern in neural tissues, we examined the 3′UTR length of the *ELAVL1* gene across 16 normal human tissues collected from the Illumina Human BodyMap 2.0 dataset (Gene Expression Omnibus, accession number GSE30611). We observed a significant 3’UTR lengthening of *ELAVL1* in brain samples compared to other tissues (**Fig. 2F**). This confirms that the 3′UTR lengthening pattern is highly neural-related, and the neuroendocrine biological processes in SCNC samples exhibit conservation with the nerve system.

### Characterization of 3′UTR lengthening in SCNC at single-cell level

To gain deeper insights into the lengthening patterns and validate the key markers identified by our model, we performed comprehensive APA profiling on 48 single-cell lung cancer samples^25^ and 6 single-cell prostate cancer samples^26^. Only epithelial cells were extracted for analysis to focus on cancer-intrinsic alterations. To estimate PAS usage at single-cell resolution while maintaining consistency with bulk-level APA analysis, we first employed SCAPE^30^ for single-cell APA quantification. Next, the PAS counts of each transcript were collapsed based on the DaPars-inferred proximal PASs to calculate the proportion of transcripts with distal PAS usage.

In the lung cancer dataset (51,364 SCNC and 3,121 non-SCNC cells), we specifically examined the lengthening APA events detected at the bulk level (**Fig. 3A**). Consistent with the high feature importance observed in our prediction model, *ELAVL1* and *PSME4* exhibited the highest fold change in pseudo-bulk PDUI, showing significant enrichment of lengthening events in SCNC cells compared to non-SCNC cells (FDR = 0.014 and 0.031, respectively; **Fig. 3B**). Additionally, the copy number variation (CNV) analysis further confirmed higher genomic instability in SCNC cells (**Supp. Fig 4**). The elevated frequency and amplitude of CNVs in SCLC genomes may provide a molecular context for the observed 3’UTR lengthening pattern, which could contribute to the malignant phenotype.

**Figure 3.**
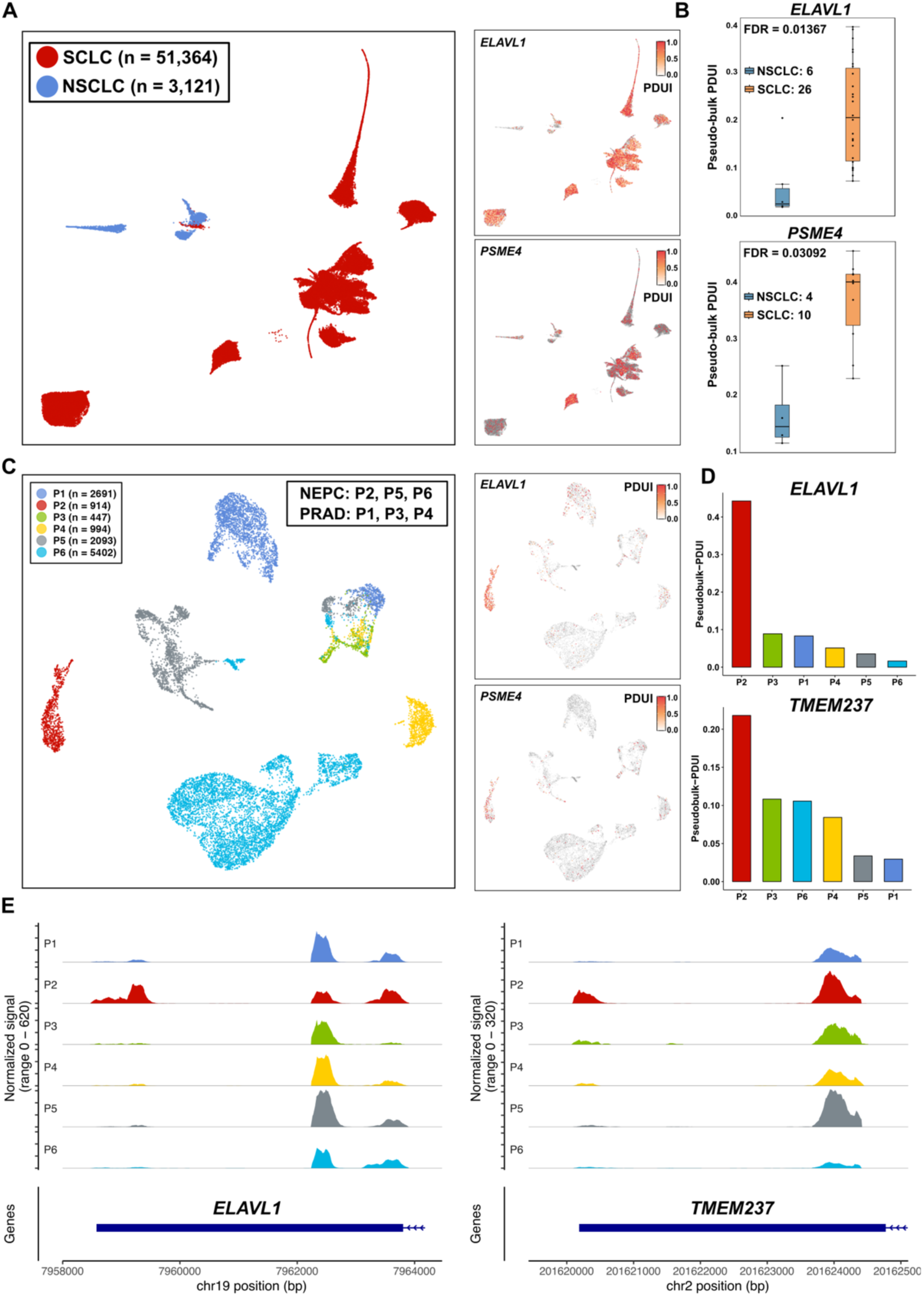
3’UTR lengthening in SCNC at single cell level. **(A)** UMAP visualization of epithelial cells from single cell lung cancer datasets. 51,364 epithelial cells of 21 SCLCs and 3,121 epithelial cells of 24 NSCLCs were extracted for visualization. The two lengthening APA events, *ELAVL1* and *PSME4*, exhibiting the highest fold change in PDUI at the pseudo-bulk level were utilized to generate the feature plots. **(B)** Boxplots comparing pseudo-bulk PDUI values of *ELAVL1* and PSME4 between NSCLC (n=6) and SCLC (n=26) samples. SCLC exhibits significantly higher PDUI values indicating increased distal PAS usage in *ELAVL1* and PSME4 for SCNC, with FDR equal to 0.01367 and 0.03092 separately. **(C)** UMAP visualization of epithelial cells from single cell prostate cancer datasets. 8,409 epithelial cells of 3 NEPC and 4,132 epithelial cells of 3 non-NEPC samples were extracted for visualization. The two lengthening APA events, *ELAVL1* and *TMEM237*, exhibiting the highest fold change in PDUI at the pseudo-bulk level were utilized to generate the feature plots. **(D)** Histograms comparing pseudo-bulk PDUI values of *ELAVL1* and *TMEM237* at the patient level. Both showed a higher pseudo-bulk PDUI level in patient #P2, but not in the other two NEPC patients. **(E)** Coverage plot of *ELAVL1* and *TMEM237* in 6 prostate cancer samples at the patient level. Only patient #2 exhibited significant distal 3’UTR coverage.

For the prostate cancer samples (8,409 SCNC and 4,132 non-SCNC cells), patients #2, #5 and #6 were clinically classified as NEPC, while patient #1, #3 and #4 were classified as non-NEPCs^26^ (**Fig. 3C**). Again, the two tumors with the highest fold change in pseudo-bulk PDUI, *ELALV1* and *TMEM237*, exhibited enrichment of lengthening APA events in patient #2. However, in the other 2 NEPC samples (patient #5 and #6), *ELAVL1* and *TMEM237* showed a low pseudo-bulk PDUI value (less than 0.1) (**Fig. 3D**), indicating only minor differences between these NEPC samples and the non-NEPC samples at the APA level. The 3′UTR coverage plot also showed the preference for the distal 3′UTR of two example APA events in patient #2 (**Fig. 3E**). We consider this could be related to the data sparsity in the APA matrix or the incomplete transdifferentiation of adenocarcinoma to neuroendocrine carcinoma that occurs during AR-directed treatment resistance. After imputing the substantial missing values with the mean PDUI value across all cells, the overall PDUI distribution may become more homogeneous, potentially obscuring less pronounced differences. Overall, these results demonstrate that APA analysis is effective in detecting significant differences between SCNC and non-SCNC samples. However, the sensitivity of this approach when used in isolation appears to be limited.

### SCNC Subtype Prediction in the CCLE Dataset

To further validate the robustness of the prediction model on a large-scale dataset, we applied it to categorize 826 cancer cell lines from the Cancer Cell Line Encyclopedia project (CCLE)^22, 23^. Using CCLE tumor subtype annotations as ground truth, our model classified 812 cell lines as non-SCNCs and 14 as SCNCs. Specifically, 11 of the 14 predicted SCNC cell lines are neuroblastomas, a neural-related tumor type. The other 3 cell lines originated from stomach, thyroid and breast cancer, respectively (**Fig. 4A**). The stomach cancer cell line *ECC10-STOMACH* and the thyroid cancer cell line *TT-THYROID* have been annotated as small-cell cancer cell lines^31, 32^, while the breast cancer cell line *MDAMB134VI_BREAST* was annotated as a non-small cell cancer cell line.

**Figure 4.**
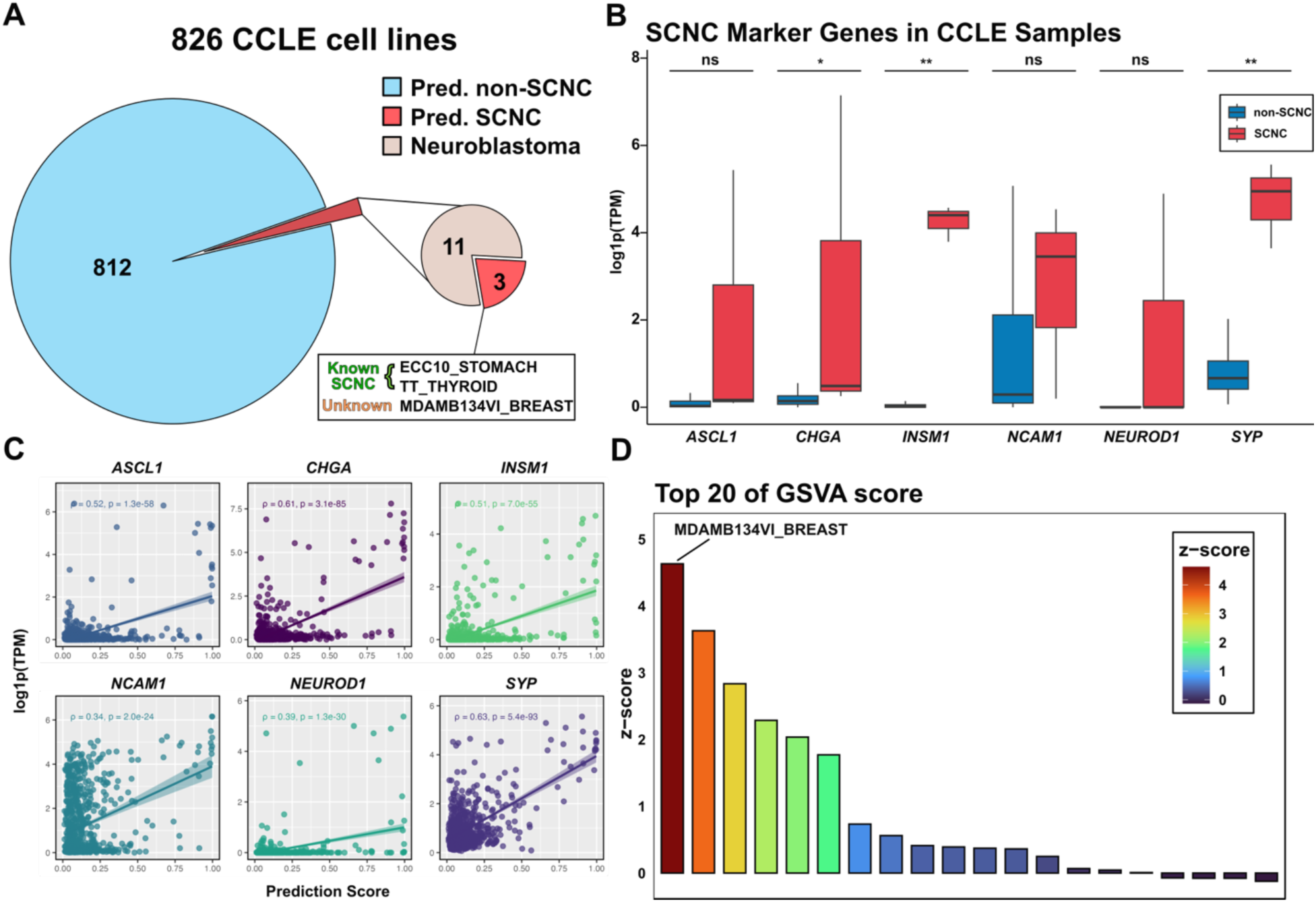
SCNC Subtype Prediction in CCLE Dataset. **(A)** Prediction results for 826 CCLE cancer cell lines: 14 samples were classified as SCNC, including 11 neuroblastoma lines, 2 annotated small-cell cancer cell lines (ECC10_STOMACH, TT_THYROID), and 1 breast cancer cell line MDAMB134VI_BREAST without prior SCNC annotation. **(B)** Expression levels (log1p(TPM)) of 6 SCNC marker genes in SCNC vs. non-SCNC cell lines. *CHGA*, *INSM1*, and *SYP* showed significant upregulation in SCNC lines (Wilcoxon rank-sum test, *p<0.05, **p<0.01). **(C)** Correlation between expression of 6 SCNC marker genes and prediction scores, showing positive associations. **(D)** Top 20 z-scores for 51 breast cancer cell lines based on *CHGA*, *INSM1*, and *SYP* expression. MDAMB134VI_BREAST exhibits the highest z-score, suggesting potential SCNC characteristics.

To further characterize these predictions, we quantified the expression levels of six canonical SCNC marker genes (*ASCL1, CHGA, INSM1, NCAM1, NEUROD1, and SYP*). We compared the predicted SCNC cell lines (excluding neuroblastoma) against all non-SCNC cell lines at the gene level using the Wilcoxon rank-sum test. The gene *CHGA, INSM1, and SYP* exhibited significant upregulation in the three predicted SCNC cell lines, whereas differences in the other three markers were not statistically significant (**Fig. 4B**). Previous studies on SCNC and non-SCNC classification have noted that individual tumors may exhibit high expression of only a subset of SCNC marker genes, potentially leading to misclassification when using traditional schemes that rely on just a few markers^5^. Our findings align with this observation, reinforcing the importance of comprehensive marker profiling for tumor subtype classification.

Additionally, we evaluated the correlation between the gene expression level of each SCNC marker gene and the prediction score across samples. With the exception for *NCAM1* and *NEUROD1,* prediction scores exhibited a moderate positive correlation with the expression level of the other 4 SCNC marker genes, supporting the prediction results (**Fig. 4C, Supp. Fig 5**). Focusing on the markers significantly upregulated in the predicted SCNC group (*CHGA*, *INSM1*, and *SYP*), we examined their expression across 51 breast cancer cell lines to investigate the status of MDAMB134VI_BREAST to assess whether it represents a potential SCNC cell line. Using Gene Set Variation Analysis (GSVA) to calculate the composite z-scores based on these three markers, we found that *MDAMB134VI_BREAST* ranked at the top among all the breast cancer cell lines (**Fig. 4D**), indicating the highest expression levels of these three SCNC marker genes in this sample and a high probability of it being a SCNC cell line.

### SCNC Subtype Prediction in the TCGA Datasets

Following the prediction validation on datasets with known tumor subtypes, we applied the APA signature model to identify potential SCNC samples among 7,659 tumors from TCGA datasets. Of these, 7,600 samples were classified as non-SCNCs, and 59 samples were classified as SCNCs. Specifically, 28 of the predicted SCNC samples are glioblastoma (GBM), a neural-related tumor type, and the remaining 31 samples belong to other cancer types (**Fig. 5A**). Similar to the validation on the CCLE datasets, we quantified the gene expression level of *ASCL1, CHGA, INSM1, NCAM1, NEUROD1, and SYP* to compare the SCNC (excluding GBM) and non-SCNC samples at the gene level using the Wilcoxon rank-sum test. With the exception of *INSM1,* all the other markers were significantly upregulated in the 31 predicted SCNC samples (**Fig. 5B**), strongly suggesting that these samples possess SCNC characteristics.

**Figure 5.**
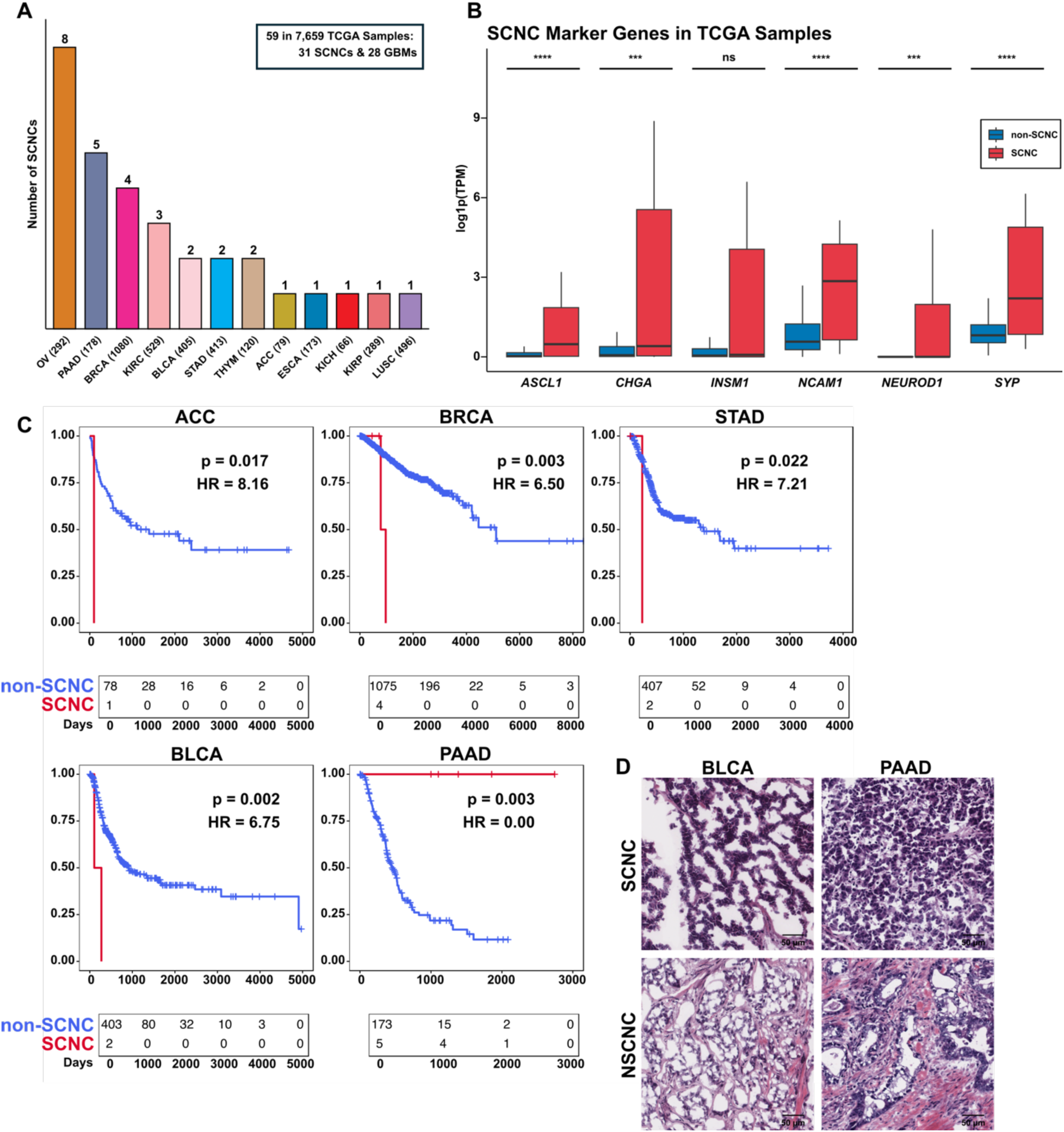
SCNC Subtype Prediction in TCGA Dataset. **(A)** Prediction results for 7,659 pan-cancer samples from TCGA: 59 classified as SCNCs, including 28 glioblastomas and other 31 samples of 12 cancer types. **(B)** Expression levels (log1p(TPM)) of 6 SCNC marker genes in SCNC vs. non-SCNC samples. Except for INSM1, the other 5 marker genes showed significant upregulation in SCNC samples (Wilcoxon rank-sum test, *p<0.05, **p<0.01, ***p<0.001, ****p<0.0001). **(C)** Progression-free interval survival analysis of predicted SCNC samples. Significant outcome differences observed in 5 cancer types: poorer survival in SCNCs of ACC, BRCA, BLCA, and STAD; better survival in SCNCs of PAAD. **(D)** Representative pathology slides from flash frozen samples: SCNC of BLCA (TCGA-G2-A2EL, top left), SCNC of PAAD (TCGA-3A-A9IL, top right), non-SCNC of BLCA (TCGA-FD-A3SN, bottom left), and non-SCNC of PAAD (TCGA-2J-AABT, bottom right). Scale bar: 50 µm.

To further characterize the clinical relevance of these predictions, we performed survival analysis across 12 tumor types containing potential SCNC cases to investigate prognostic differences between SCNC and non-SCNC groups. Strikingly, the predicted SCNC samples in 5 tumor types exhibited significant prognostic disparities in Progression-free survival interval (PFI), indicating the capability of this model to identify previously unrecognized SCNC cases. In adrenocortical carcinoma (ACC), breast invasive carcinoma (BRCA), bladder urothelial carcinoma (BLCA), and stomach adenocarcinoma (STAD), SCNC samples were associated with significantly worse survival outcomes than their non-SCNC counterparts (Hazard Ratio > 6.50, p < 0.05 for all). Conversely, SCNC samples in pancreatic adenocarcinoma (PAAD) showed markedly improved survival (Hazard Ratio = 0.00, p = 0.003) (**Fig. 5C**), consistent with reports in the literature for pancreatic neuroendocrine studies^33^.

Furthermore, we examined the available TCGA hematoxylin and eosin (H&E) stained histology slides and pathological reports available at the Cancer Digital Slide Archive^34^. The 7 predicted SCNC samples of BLCA and PAAD were explicitly annotated as SCNC samples or noted to have extensive small-cell differentiation in the pathology reports. Representative pathological slides of validated SCNC and non-SCNC samples are listed (**Fig. 5D**). The confirmed SCNC cells exhibited classic morphological features, including high nuclear-to-cytoplasm ratios, nuclear molding, and stippled chromatin. Specifically, a sample of lung squamous cell carcinoma (LUSC) was also predicted as SCNC. To validate our findings, we leveraged differential gene expression patterns between SCLC and NSCLC, previously identified from 192 bulk RNA-Seq samples We then applied the Virtual Inference of Protein-activity by Enriched Regulon analysis (VIPER) algorithm^35^ to score the samples of LUSC based on this differential gene set. Interestingly, the sample predicted as SCNC by our model exhibited the highest VIPER scores (**Supp. Fig 6**), suggesting the potential presence of unannotated SCNC samples within the TCGA-LUSC cohort.

### Differential lengthening of ELAVL1 transcripts in small-cell lung cancer cell lines

To validate the identified 3′UTR lengthening pattern in SCNC, we focused on *ELAVL1*, one of the most significant lengthening genes from our previous results. We first examined the 3′UTR of *ELAVL1* in six human lung cancer cell lines: three human NSCLC cell lines (H1299, H460, A549) and three human SCLC cell lines (H524, H2171, H1963). The coverage plot of *ELAVL1* in the six cell lines validated a strong preference for the distal PAS of 3’UTR of *ELAVL1* (**Fig. 6A**). Next, we performed RT-PCR analysis to quantify the ratio of long 3’UTR to total ELAVL1 mRNA across three human NSCLC cell lines (H1299, H460, A549) and the three human SCLC cell lines (H524, H2171, H1963) (**Fig. 6B**). The results showed a striking difference between NSCLC and SCLC cell lines. In NSCLC cell lines (H1299, H460, A549), the long 3′UTR/total ratio was consistently low, with values of 0.025, 0.002, and 0.023, respectively. In contrast, SCLC cell lines (H524, H271, H1963) exhibited significantly higher ratios of 0.446, 0.398, and 0.611, respectively. These data strongly support a substantial increase in both the proportion of ELAVL1 transcripts with longer 3′UTRs and the alternative isoforms in SCLC compared to NSCLC cell lines.

**Figure 6.**
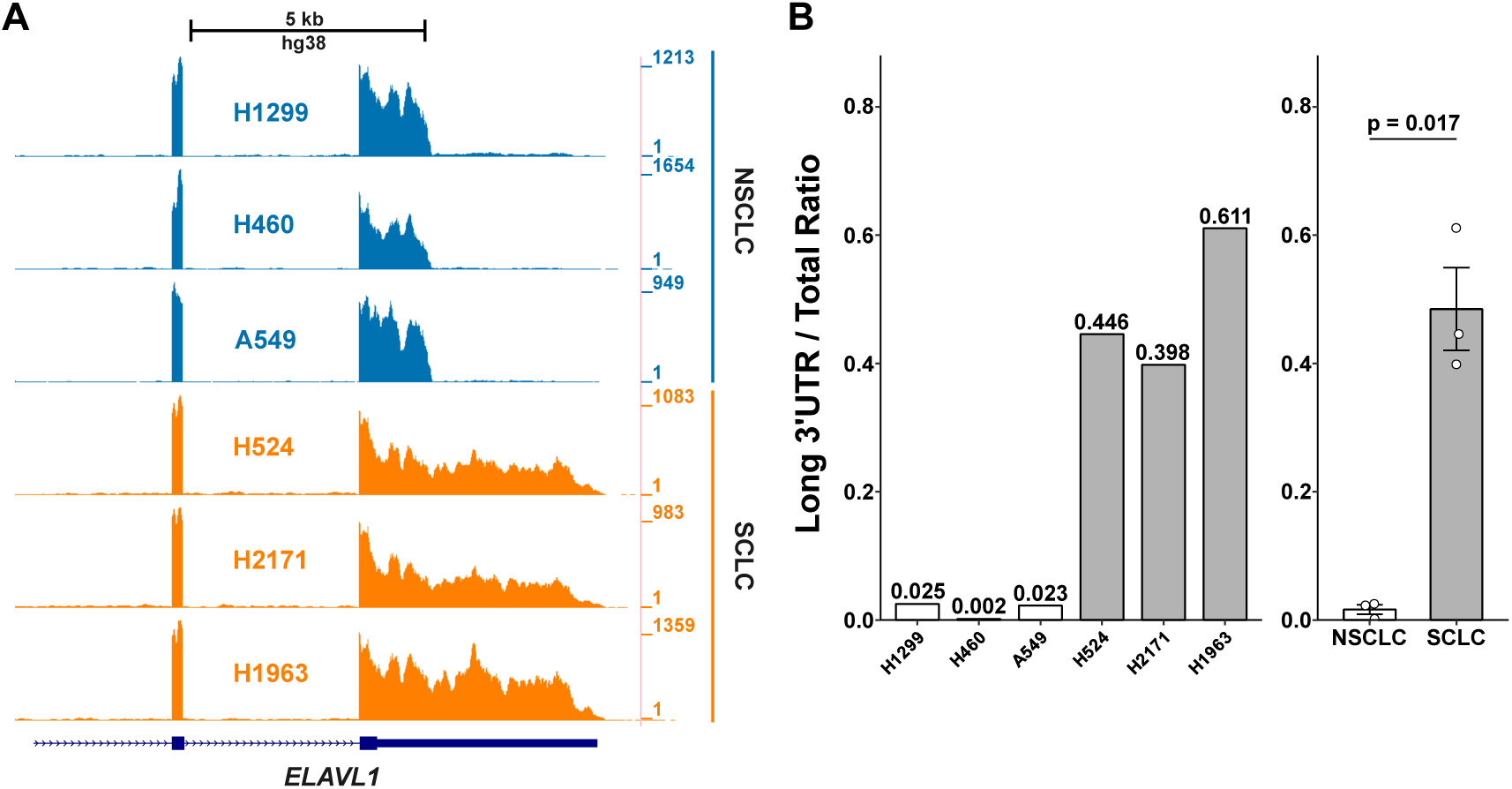
ELAVL1 Lengthened in SCLC cell lines. **(A)** Coverage plots of ELAVL1 in 3 NSCLC and 3 SCLC cell lines. ELAVL1 showed a longer mRNA isoforms preference in the SCLC cell lines. **(B)** Results of RT-PCR of ELAVL1. A high long 3’UTR ratio was observed in the SCLC cell lines.

## Discussion

In this study, we characterized the landscape of alternative polyadenylation (APA) within SCNC and non-SCNC samples across multiple tissues. We revealed a previously unrecognized 3’UTR length pattern that distinguishes these tumor subtypes through quantification of PAS usage and an APA signature-based prediction model. While numerous APA-related studies have focused on dynamic PAS usage between tumor and normal tissues^36–38^, the landscape of APA among different tumor subtypes has remained largely unexplored.

The conventional paradigm holds that transcripts with shorter 3’UTRs are preferentially expressed in tumors^17^. This preference arises from reduced miRNA binding sites in truncated 3’UTRs, allowing these transcripts to evade miRNA-mediated degradation^16, 39^. However, we uncovered a global mRNA 3’UTR lengthening pattern in SCNC, which are typically more malignant than non-SCNC. This paradoxical finding was validated across both lung and prostate SCNC samples, with a conserved set of lengthening APA signatures identified. This counterintuitive observation suggests that SCNC may employ fundamentally different post-transcriptional regulatory mechanisms compared to conventional carcinomas. It potentially reflects the activation of neural-specific gene expression programs rather than simply representing dysregulated APA processing.

Traditional approaches to identifying SCNC have relied on neuroendocrine marker genes such as synaptophysin (SYP) and chromogranin A (CHGA)^5^. Our study extends this molecular toolkit by leveraging SCNC-specific APA events resulting in 3’UTR lengthening in genes such as ELAVL1, as complementary molecular markers for tumor classification. The consistency between APA-based classification and traditional marker gene expression levels demonstrates that post-transcriptional features can serve as orthogonal biomarkers. To operationalize this finding, we developed a neural network model utilizing 20 conserved APA lengthening signatures to predict potential SCNC samples. This provides a systematic framework for SCNC identification at the post-transcriptional level.

Our analysis integrated both bulk and single-cell RNA-seq data to obtain comprehensive insight into APA patterns. From tissues commonly associated with SCNC, we profiled 49 prostate cancer^21^ (15 NEPCs & 34 PRADs) and 192 lung cancers^22, 23^ (50 SCLCs & 142 NSCLCs) samples at the bulk RNA-Seq level using DaPars^40^. We identified hundreds of APA events with distal PAS preference in SCNC versus only approximately 10 shortened events. To ensure consistency between platforms and mitigate technical biases, we utilized PASs identified by DaPars in bulk RNA-seq as references for single-cell analysis. We applied SCAPE^30^ across 45 lung cancer^25^ and 6 prostate cancer^26^ samples. This integration strategy confirmed that bulk-level lengthening events, such as ELAVL1, exhibited enrichment in SCNC cells with significant pseudo-bulk PDUI differences between SCNC and non-SCNC samples.

Intriguingly, the lengthening APA signatures showed similar distal PAS preference in normal brain tissue compared to 15 other normal human tissues. These signatures were enriched in brain-related pathways. This molecular convergence between SCNC and neural tissue provides a mechanistic basis for the neural-like characteristics of these tumors. It suggests that neuroendocrine differentiation may recapitulate developmental programs active in brain tissue. The functional consequences of this recapitulation warrant further investigation. Specifically, we need to determine whether 3’UTR lengthening confers selective advantages through gain of regulatory elements, altered mRNA localization, or other post-transcriptional mechanisms. Understanding the regulatory factors mediating this APA shift represents a critical avenue for future mechanistic studies. Core cleavage and polyadenylation machinery components, such as the CFIm complex, likely play pivotal roles.

However, heterogeneity among samples suggests that APA analysis alone may have limitations. Among NEPC samples, patient #2 exhibited pronounced 3’UTR lengthening, while patients #5 and #6 showed no significant differences compared to non-NEPC samples. This variability indicates that combining APA analysis with other analytical methods could improve sensitivity and comprehensiveness in differentiating SCNC from non-SCNC. Such integration may reveal subtle variations that might be obscured when relying solely on APA data. Multi-omic approaches may be particularly valuable for capturing the spectrum of neuroendocrine differentiation states. To establish robust and generalizable signatures, we analyzed PDUI levels across over 7,600 tumor samples from TCGA cohorts. We identified 20 conserved lengthening APA signatures that served as inputs for our neural network classification model. Validation using 6 prostate cancer single-cell RNA-seq datasets and 826 CCLE cancer cell lines with subtype annotations confirmed the model’s performance and minimized overfitting concerns. The model successfully classified prostate cancer patient #2 as SCNC at the pseudo-bulk level while correctly labeling the remaining 5 samples as non-SCNC, consistent with APA profiling results. Among three predicted SCNC cell lines from CCLE, two were confirmed SCNC types. The third—a breast cancer cell line—exhibited high expression of SYP and INSM1 with top-ranked z-scores for SCNC markers (CHGA, INSM1, SYP). This suggests it may represent an unannotated SCNC case. Application of the model to TCGA cohorts predicted 31 potential SCNC samples. These samples demonstrated significant upregulation of traditional SCNC marker genes and distinct clinical outcomes in ACC, BLCA, BRCA, PAAD, and STAD. Pathology slides and histological reports further verified predictions in BLCA and PAAD, demonstrating the practical utility of APA signatures for SCNC identification. These validations collectively establish that APA patterns can effectively serve as additional molecular markers with significant discriminatory power.

Several limitations should be acknowledged. The sample sizes for some analyses, particularly single-cell datasets, remain modest and may limit statistical power for detecting subtle effects. While we validated predictions using available clinical annotations and pathology reports, prospective validation in independent cohorts will be essential for clinical translation. Functionally, whether 3’UTR lengthening alters protein abundance or function, actively drives SCNC pathogenesis, or primarily serves as a biomarker remains to be determined. Targeted experimental approaches such as 3’UTR reporter assays, CRISPR-based PAS manipulation, and functional genomics screens will be needed. Additionally, investigating whether APA patterns correlate with therapeutic response could establish clinical utility beyond diagnostics.

In conclusion, our study uncovers a distinct mRNA 3’UTR lengthening pattern in SCNC and demonstrates the utility of APA signatures for cancer subtype classification. The integration of APA analysis with traditional gene expression markers and machine learning techniques offers a multi-dimensional approach to understanding tumor biology. As we continue to refine and validate this model across diverse cohorts, it has the potential to become a valuable tool in personalized cancer diagnostics and treatment planning. Future investigations should prioritize three key areas: elucidating the functional consequences of 3’UTR lengthening in SCNC, exploring how this phenomenon contributes to tumor aggressiveness, and identifying the regulatory mechanisms mediating APA changes. These efforts could uncover new therapeutic targets for managing these challenging malignancies.

## Methods

### Sample Description

The bulk RNA-Seq data of prostate cancer including 49 prostate cancer cases^21^ from dbGaP under accession phs000909, and 192 lung cancer cell lines from CCLE^22, 23^. The data of four patient-derived xenografts (PDXs) prostate cancer samples (2 NEPCs and 2 PRADs) from the European Nucleotide Archive under accession number ENA: PRJEB9660^24^. The data of normal human tissues were collected from the Illumina Human BodyMap 2.0 dataset (Gene Expression Omnibus accession code GSE30611). For APA characterizing across tissues, the data from other cancer types, including 826 tumor samples from CCLE and 7,659 tumor samples from TCGA were also incorporated. For single-cell RNA-Seq data, 6 prostate cancer samples and 45 lung cancer samples were collected from Gene Expression Omnibus (GEO) under accession GSE137829^26^ and dbGaP under accession phs002371^25^. In addition, the histology reports and pathology slides of the TCGA samples have been collected from the Cancer Digital Slides Archive (CDSA)^34^ to validate the APA-based SCNC classification and prediction results.

### Bulk RNA-Seq Data Processing

The raw fastq files were first quality checked using FastQC (version 0.11.8). Raw fastq files were aligned to hg38 human reference genome (GRCh38.84) via HISAT2^41^ (v2.2.1) and per-gene counts quantified by RSEM^42^ (version 1.3.1) based on the gene annotation Homo_sapiens.GRCh38.84.chr.gtf. Raw read counts were processed and normalized to transcripts per million (TPM) using the RSEM algorithm (v1.3.1). Post-alignment processing was conducted using SAMtools^43^ (version 1.20) to sort and index aligned reads, followed by extraction of reads mapping to chromosomes 1-22, X, Y, and MT. The resulting BAM files were then re-indexed, and chromosome size information was extracted from the processed files. To generate genome coverage data, the *genomeCoverageBed* function from BEDTools^44^ (v2.31.1) was applied to convert the aligned BAM files to BedGraph format with the *sort* parameter to sort the BedGraph files. The bedGraphToBigWig utility from the UCSC Genome Browser toolkit^45^ was utilized to convert the sorted BedGraph files to bigWig format for visualization in genome browser. Differential gene expression values were obtained using DESeq2^46^ (version 1.42.1). Gene expression differences were considered significant if they passed the following criteria: adjusted P value < 0.05, |log2(fold change)| >= log2(1.5).

### Single-cell RNA-Seq Data Processing

The sequencing data were processed using CellRanger software (10x Genomics Cell Ranger v7.1.0) with default parameters and mapped to the human genome (GRCh38.84). Subsequent analyses were performed using the Seurat package^47^ (version 5.1.0) in R. The low-quality cells (gene count < 200 or the mitochondrial gene ratio > 10%) were removed. The normalization was performed using SCTransform^48^ with default parameters. Following normalization, dimensionality reduction was performed using Principal Component Analysis (PCA) on the top 2000 highly variable genes. The first 30 principal components were used for downstream analyses. Uniform Manifold Approximation and Projection (UMAP) was applied for visualization of the high-dimensional data in two-dimensional space. Clustering was performed using the Louvain algorithm with a resolution parameter of 0.5. This resulted in the identification of distinct cell clusters, which were then annotated based on the cell annotation from original studies.

### Bulk level APA Analysis

Bulk level APA analysis was performed using DaPars^40^ to estimate the PAS usage of transcripts in each sample and calculate the PDUI value represents the proportion of transcript isoforms with distal PAS. The BedGraph files of each sample generated from the bulk data processing step and the chromosome size file were utilized as the input files. Extracting the distal polyadenylation sites from the genome annotation, the proximal PAS would be inferred and utilized to quantify the PDUI. The PDUI matrix is further applied to downstream differential APA analysis and the training of prediction model.

### Single-cell level APA Analysis

Single-cell level APA analysis was performed using SCAPE^30^ to estimate the PAS usage of transcripts in each cell. With the aligned bam files and the gene expression level matrix generated by CellRanger from the scRNA-Seq data processing step, SCAPE generates a PAS counts matrix for each PAS in each cell. To mitigate batch effects and ensure consistency in identified PAS locations between bulk and single-cell datasets, we aggregated the SCAPE-inferred PASs based on the PAS locations calculated by DaPars. Using the proximal PAS identified by DaPars as a guide, the SCAPE-estimated PASs were divided into two groups, representing the proximal and distal regions of the 3’UTR. For the positive strand, the expression level of SCAPE-estimated PASs on the right side of the DaPars-identified locus were aggregated to compute the mean counts represent distal PAS usage. The mean counts of the PASs on the left side of the DaPars-identified locus represented proximal PAS usage. For the negative strand, the direction was reversed, and the same aggregation was performed to obtain the PAS usages. In this way, we calculated the PDUI value of each APA event in each single cell with consistent approach as bulk level APA analysis.

### Differential APA Analysis

At the bulk level, we utilized DaPars to characterize the APA events within 49 prostate cancer samples from dbGaP and 192 lung cancer samples from CCLE. The differential APA analysis were applied to identify the SCNC-specific APA events. In detail, Fisher’s exact test was performed to determine the p-value of PDUI difference between SCNC and non-SCNC samples, which was further adjusted by the Benjamini–Hochberg procedure to control the false-discovery rate (FDR) at 0.1. The absolute mean difference of PDUIs of all the samples in the comparison of SCNC and non-SCNC must be greater than 0.05. Meanwhile, the mean fold-change of PDUIs of all the samples in the comparison of SCNC and non-SCNC must be greater than 1.4. Further, the mean PDUI for SCNC and non-SCNC samples must exceed 0.15 to filter out APA events with deficient expression levels.

### Construction of Neuro-network-based prediction model

The neural network architecture was implemented using the *Keras* package^49^ (version 2.15.0). It comprised two dense layers and one output layer, adhering to the default configuration. The model commenced by defining a dense layer with 64 neurons and a *ReLU*^50^ activation function while specifying the input shape based on the 20 conserved APA signatures, representing the features. The missing values within the matrix of each APA event were imputed using the mean value among all samples in the matrix. The data then propagated into another dense layer with 32 neurons, maintaining the use of *ReLU* activation. The output layer was established with a single neuron and a *sigmoid*^51^ activation function, which is suitable for binary classification tasks as it yields a probability score between 0 and 1. Training of the model employed the *Adam*^52^ optimizer alongside a *binary_crossentropy* loss function across 50 epochs, utilizing a batch size of 16. The resulting prediction model generated a table containing the anticipated probabilities of samples being classified as either “SCNC” or “non-SCNC” based on the model’s predictions. Subsequently, the pseudo-bulk PDUI matrix derived from the 20 × 28 single-cell datasets of tumor-subtype-annotated lung cancer samples from dbGaP was utilized to perform testing to determine the optimal cutoff for probabilities for binary classification, conducting ROC curve analysis. Further, 10-fold cross-validation was performed on the 20 × 241 PDUI matrix from 49 prostate cancer and 192 lung cancer samples to ensure robustness and reliability in classification performance assessment. The cross-validation was implemented using the built-in functionality of *Keras* along with base R functions for data splitting. Each fold involved a random division of the dataset into 90% training and 10% testing subsets, ensuring representative training and testing sets.

### Survival Analysis

Survival analysis was conducted utilizing the R packages *survminer* (version 0.4.9) and *survival* (version 3.7-0) packages. Progression-free interval (PFI) data were extracted from The Cancer Genome Atlas (TCGA) clinical data resource. Samples were categorized as “SCNC” or “non-SCNC” based on the predictions generated by our model. For each cancer type where small cell samples were predicted, Kaplan-Meier survival curves were fitted using the *survfit* function. Hazard ratios (HR) and their corresponding 95% confidence intervals were calculated through the application of Cox proportional hazards models. Statistical significance between survival curves was assessed using the log-rank test, with a threshold of p < 0.05 considered significant.

### Gene Set Variation Analysis (GSVA)

Gene Set Variation Analysis was performed using the *GSVA* package (version 1.42.0) to assess the enrichment of SCNC marker genes across breast cancer cell lines from the CCLE dataset. The expression data were log1p-transformed and normalized prior to analysis. Six key SCNC markers (*ASCL1, CHGA*, *INSM1*, *NCAM1*, *NEUROD1*, and *SYP*) were used for the analysis. GSVA was run using the z-score method with minimum gene set size of 1 and maximum of 3 to accommodate our focused gene set.

### VIPER Analysis

Virtual Inference of Protein-activity by Enriched Regulon analysis (VIPER) was performed the *viper* package (version 1.38.0). Gene expression data were obtained from The Cancer Genome Atlas (TCGA) for lung squamous cell carcinoma (LUSC) cohorts. Differentially expressed genes (DEGs) between SCNC and non-SCNC were identified using a threshold of |log2(fold change)| >= log2(1.5) and adjusted p-value < 0.05. The top 200 up-regulated and 200 down-regulated genes, ranked by fold change, were selected to construct the regulon for VIPER analysis. VIPER was executed using the rank-based enrichment method, with a minimum regulon size of 10 genes. Resulting VIPER scores were calculated for each sample, representing the enrichment of the SCNC-specific genes.

### Cell culture, antibodies and immunoblot (IB)

Human H1299, H460, A549, H524, H2171 and H1963 lung cancer cell lines were obtained from ATCC and cultured in Dulbecco’s modified Eagle’s medium (DMEM) supplemented with 10% fetal bovine serum (FBS), 50 U/ml penicillin and 0.1 mg/ml streptomycin at 37°C in a 5% CO2 humidified atmosphere. Cell lines were passaged less than 30 times for a maximum 2 months and routinely monitored for Mycoplasma contamination. Manufacturers performed authentication through short tandem repeat profiling. Cells were lysed in lysis buffer consisting of 50 mM Tris-HCl (pH 8.0), 0.5% Nonidet P-40, 1 mM EDTA, 150 mM NaCl, 1 mM phenylmethylsulfonyl fluoride (PMSF), 1 mM DTT, 1 µg/ml pepstatin A, and 1 mM leupeptin. Equal amounts of cell lysates were used for IB analysis. anti-ELALV1 (sc-5261, Santa Cruz Biotechology, RRID:AB_627770) and anti-tubulin (66240-1-Ig, Proteintech, RRID:AB_2881629) antibodies were purchased.

### Reverse transcriptase-Quantitative polymerase chain reaction (RT-qPCR)

Total RNA was isolated from cells using Qiagen RNeasy Mini Kits (Qiagen, Valencia, CA). Reverse transcriptions were performed using RevertAid RT Reverse transcription Kit (K1691, Thermo Scientific). Quantitative real-time PCR was performed on an ABI StepOneTM real-time PCR system (Applied Biosystems) using SYBR Green Mix (Thermo Fisher Scientific) as described^53^. All reactions were carried out in triplicate. Relative gene expression was calculated using the 1′Cρ method following the manufacturer’s instruction. The primers used were listed below:

## Data Availability

The publicly available datasets used in this study can be accessed from multiple sources. The RNA sequencing data used in this study were obtained from The Cancer Genome Atlas (TCGA) database (https://portal.gdc.cancer.gov/), while the Gene Expression Omnibus (GEO, https://www.ncbi.nlm.nih.gov/geo/) database contains GSE137829^26^, GSE30611, and dbGaP (https://www.ncbi.nlm.nih.gov/gap/; phs000909^21^ and phs002371^25^) is another source. The datasets from European Nucleotide Archive can be accessed under the accession number PRJNA523380^22, 23^ and PRJEB9660^24^. The author can provide any remaining data upon reasonable request.

## Code Availability

All analyses were conducted using publicly available software and packages. All custom code used in this study is available upon reasonable request from the corresponding author.

## Acknowledgements

This work was supported by the following funding: Department of Defense Idea Development Award W81XWH2110539, NIH R01GM147365 and Silver Family Innovation Foundation Award (to Z.X.). NIH R01CA262104 (to M.-S.D.). P50CA097186 R01, CA266452 P01, CA298991, P01CA163227, R01CA280056 (to PSN). NIH R01CA251245, NIH R01CA282005, NCCN/Pfizer/Astellas Award, Joint Institute for Cancer Research Award, and Prostate Cancer Foundation (to J.J.A.). The research reported here used computational infrastructure supported by the Office of Research Infrastructure Programs, Office of the Director, of the National Institutes of Health under Award Number S10OD034224. The results shown in bulk level APA analysis are in part based upon data generated by the TCGA Research Network: https://www.cancer.gov/tcga. The content is solely the responsibility of the authors and does not necessarily represent the official views of the funders.

## Author contributions

Z.X. conceived the idea. Y.Z. and Z.X. performed computational analyses. Y.Z., X.Z., Y.-M.H., F.Z., S.D. and Z.X. interpreted data. R.S.D., X.-X.S. and M.-S.D. performed experiments. H.W. and N.K.A. reviewed and interpreted the pathology images. R.C.S., E.C., J.R.B., J.J.A., G.B.M. and P.S.N. provided input on manuscript content, data interpretation, and clinical insights. Z.X. supervised the study. Y.Z. and Z.X. wrote the manuscript. All other authors provided critical feedback and approved the final manuscript.

