## Supplemental Figures for "Global mRNA 3′UTR lengthening in small-cell neuroendocrine carcinoma"

### Supp. Fig 1

#### APA in PDX

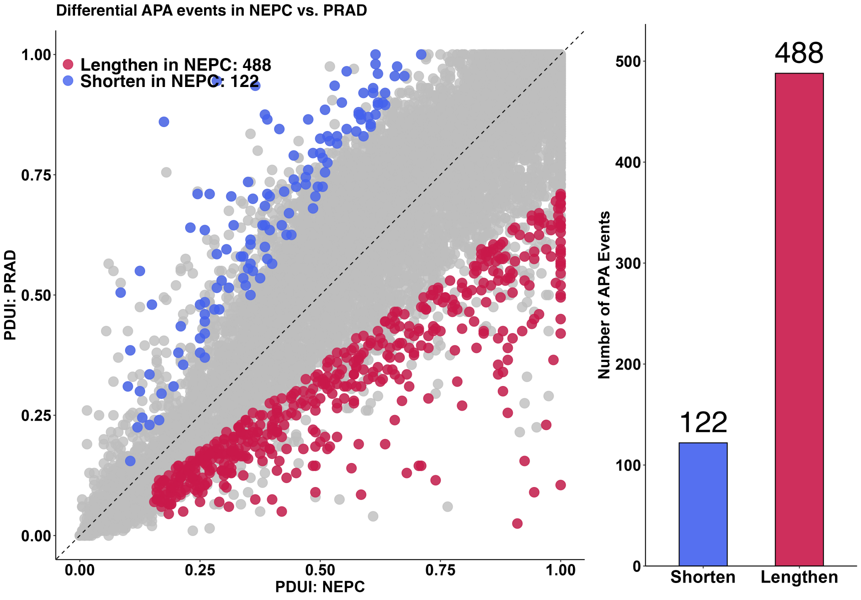


#### APA in healthy human tissues

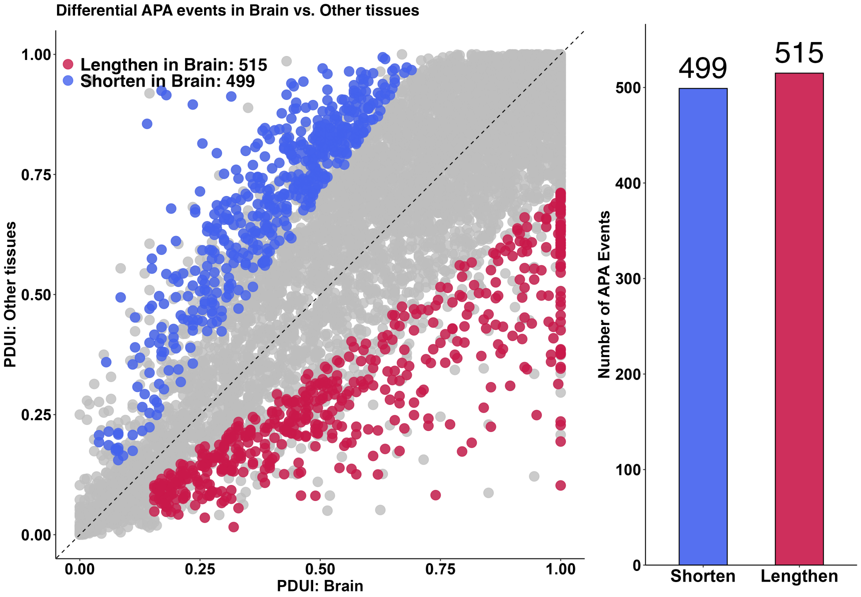

### Supp. Fig 2

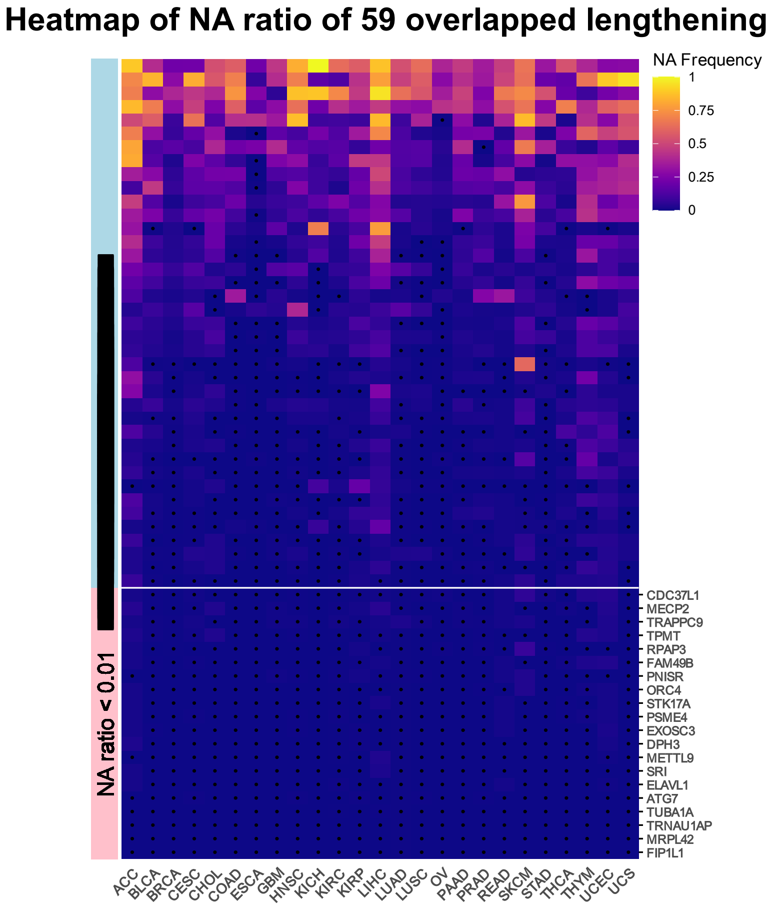

Heatmap of NA ratio of 59 overlapped lengthening APA
NA Frequency
1
0.75
0.5
0.25
0
[redacted]
NA ratio < 0.01
CDC37L1
MECP2
TRAPPC9
TPMT
RPAP3
FAM49B
PNISR
ORC4
STK17A
PSME4
EXOSC3
DPH3
METTL9
SRI
ELAVL1
ATG7
TUBA1A
TRNAU1AP
MRPL42
FIP1L1
ACC
BLCA
BRCA
CESC
CHOL
COAD
ESCA
GBM
HNSC
KICH
KIRC
KIRP
LIHC
LUAD
LUSC
OV
PAAD
PRAD
READ
SKCM
STAD
THCA
THYM
UCEC
UCS


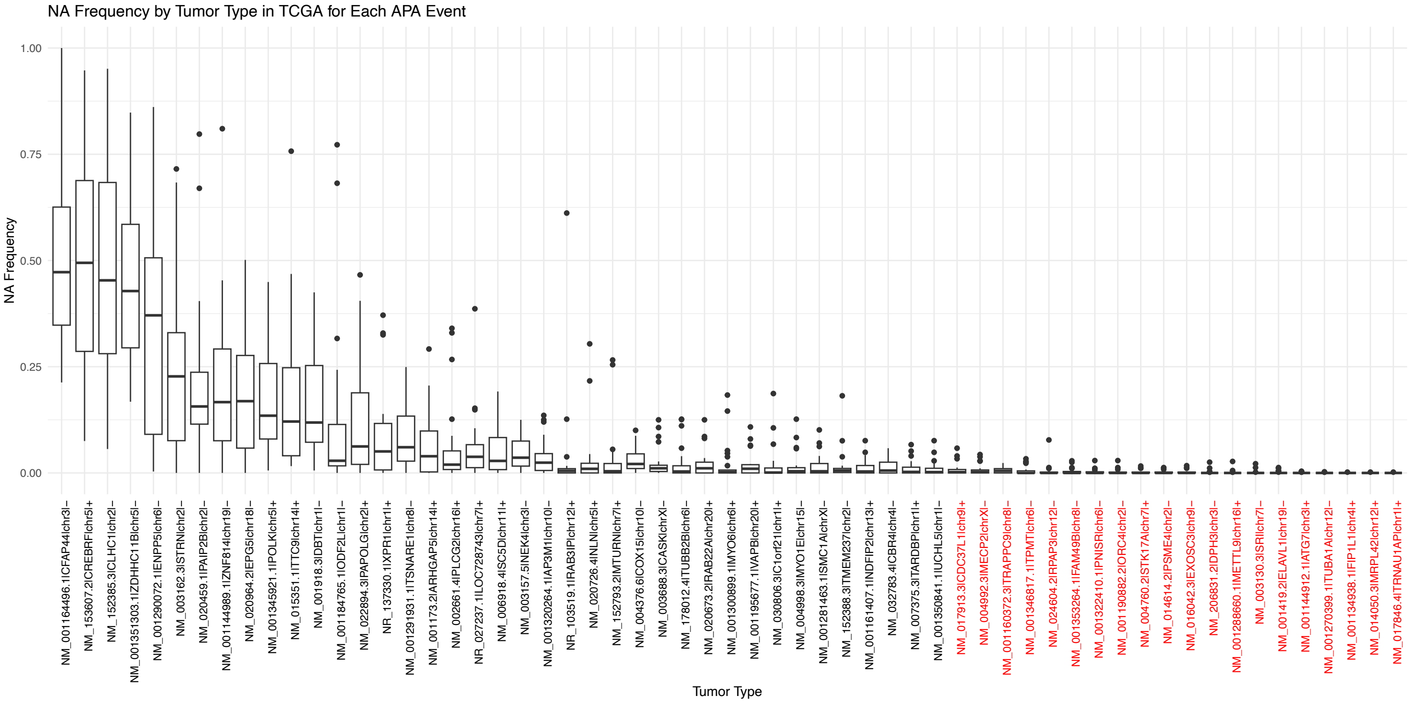

NA Frequency by Tumor Type in TCGA for Each APA Event
NA Frequency
1.00
0.75
0.50
0.25
0.00
NM_001164496.1|CFAP44|chr3|–
NM_153607.2|CREBRF|chr5|+
NM_152385.3|CLHC1|chr2|–
NM_001351303.1|ZDHHC11B|chr5|–
NM_001290072.1|ENPP5|chr6|–
NM_003162.3|STRN|chr2|–
NM_020459.1|PAIP2B|chr2|–
NM_001144989.1|ZNF814|chr19|–
NM_020964.2|EPG5|chr18|–
NM_001345921.1|POLK|chr5|+
NM_015351.1|TTC9|chr14|+
NM_001918.3|DBT|chr1|–
NM_001184765.1|ODF2L|chr1|–
NM_022894.3|PAPOLG|chr2|+
NR_137330.1|XPR1|chr1|+
NM_001291931.1|TSNARE1|chr8|–
NM_001173.2|ARHGAP5|chr14|+
NM_002661.4|PLCG2|chr16|+
NR_027237.1|LOC728743|chr7|+
NM_006918.4|SC5D|chr11|+
NM_003157.5|NEK4|chr3|–
NM_001320264.1|AP3M1|chr10|–
NR_103519.1|RAB3IP|chr12|+
NM_020726.4|NLN|chr5|+
NM_152793.2|MTURN|chr7|+
NM_004376.6|COX15|chr10|–
NM_003688.3|CASK|chrX|–
NM_178012.4|TUBB2B|chr6|–
NM_020673.2|RAB22A|chr20|+
NM_001300899.1|MYO6|chr6|+
NM_001195677.1|VAPB|chr20|+
NM_030806.3|C1orf21|chr1|+
NM_004998.3|MYO1E|chr15|–
NM_001281463.1|SMC1A|chrX|–
NM_152388.3|TMEM237|chr2|–
NM_001161407.1|NDFIP2|chr13|+
NM_032783.4|CBR4|chr4|–
NM_007375.3|TARDBP|chr1|+
NM_001350841.1|UCHL5|chr1|–
NM_017913.3|CDC37L1|chr9|+
NM_004992.3|MECP2|chrX|–
NM_001160372.3|TRAPPC9|chr8|–
NM_001346817.1|TPMT|chr6|–
NM_024604.2|RPAP3|chr12|–
NM_001353264.1|FAM49B|chr8|–
NM_001322410.1|PNISR|chr6|–
NM_001190882.2|ORC4|chr2|–
NM_004760.2|STK17A|chr7|+
NM_014614.2|PSME4|chr2|–
NM_016042.3|EXOSC3|chr9|–
NM_206831.2|DPH3|chr3|–
NM_001288660.1|METTL9|chr16|+
NM_003130.3|SRI|chr7|–
NM_001419.2|ELAVL1|chr19|–
NM_001144912.1|ATG7|chr3|+
NM_001270399.1|TUBA1A|chr12|–
NM_001134938.1|FIP1L1|chr4|+
NM_014050.3|MRPL42|chr12|+
NM_017846.4|TRNAU1AP|chr1|+
Tumor Type

**Supp. Fig 3**

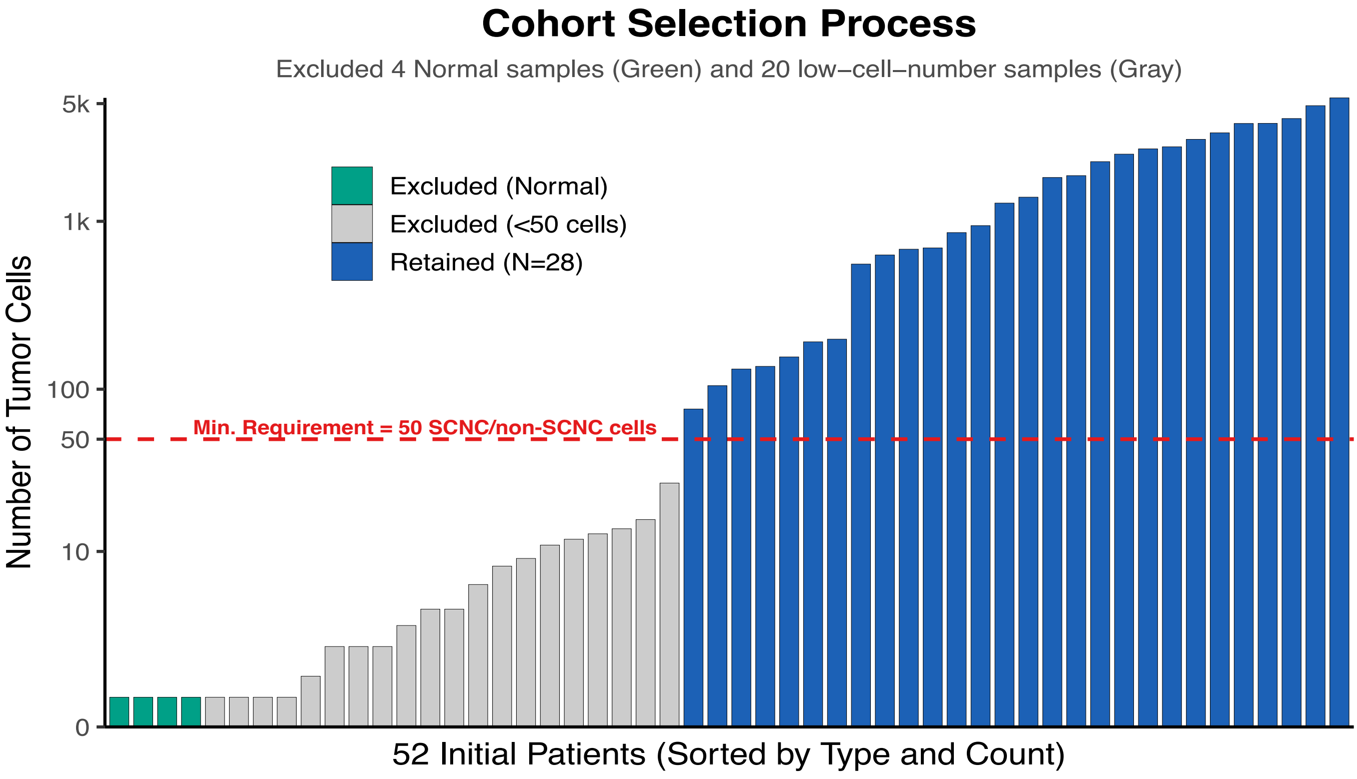
Cohort Selection Process
Excluded 4 Normal samples (Green) and 20 low–cell–number samples (Gray)
Excluded (Normal)
Excluded (<50 cells)
Retained (N=28)
Min. Requirement = 50 SCNC/non-SCNC cells
5k
1k
100
50
10
0
Number of Tumor Cells
52 Initial Patients (Sorted by Type and Count)

### Supp. Fig 4

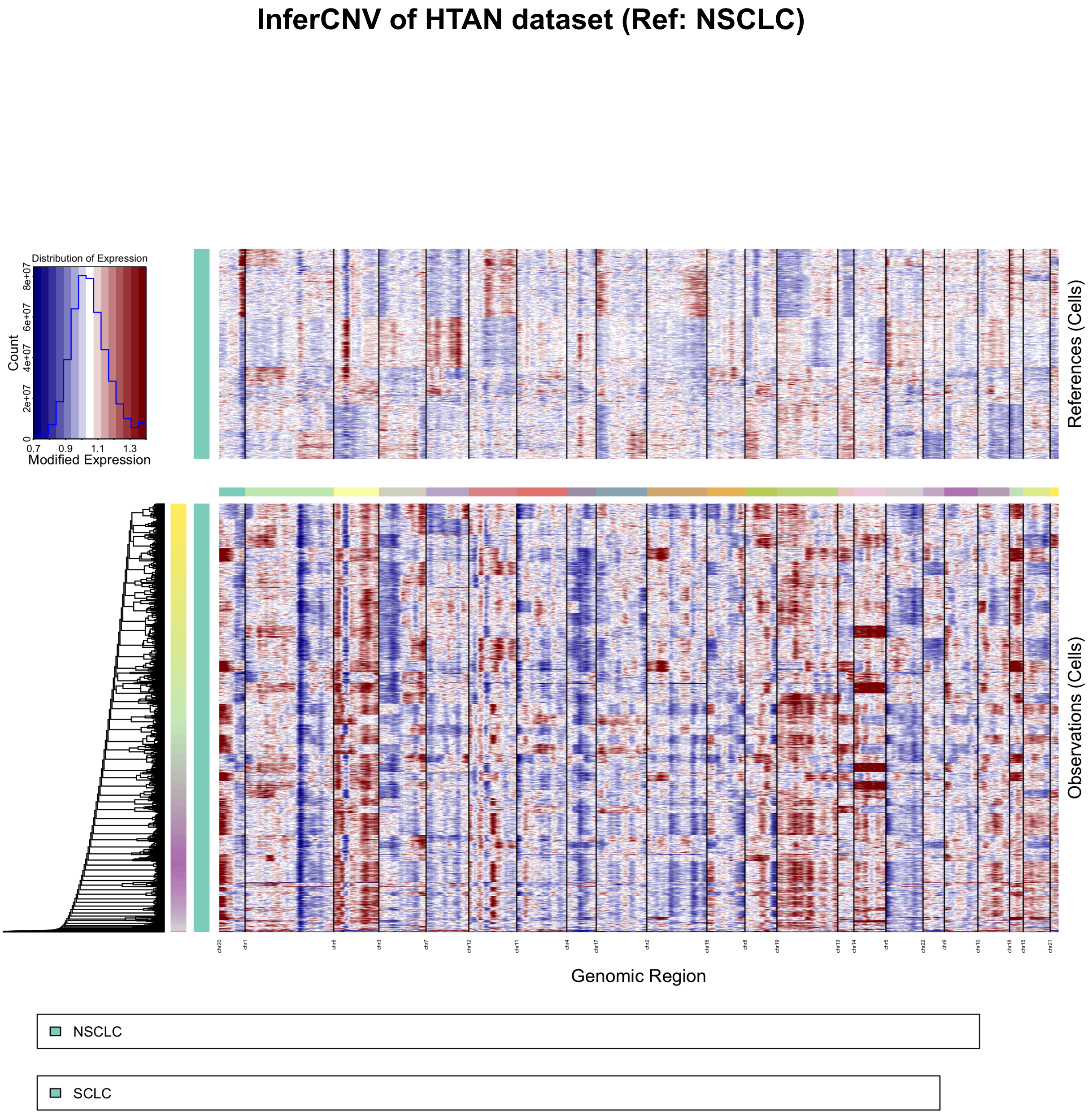

InferCNV of HTAN dataset (Ref: NSCLC)
Distribution of Expression
Count
0
2e+07
4e+07
6e+07
8e+07
0.7
0.9
1.1
1.3
Modified Expression
References (Cells)
Observations (Cells)
chr20
chr1
chr6
chr3
chr7
chr12
chr11
chr4
chr17
chr2
chr16
chr8
chr19
chr13
chr14
chr5
chr22
chr9
chr10
chr18
chr15
chr21
Genomic Region
NSCLC
SCLC

### Supp. Fig 5

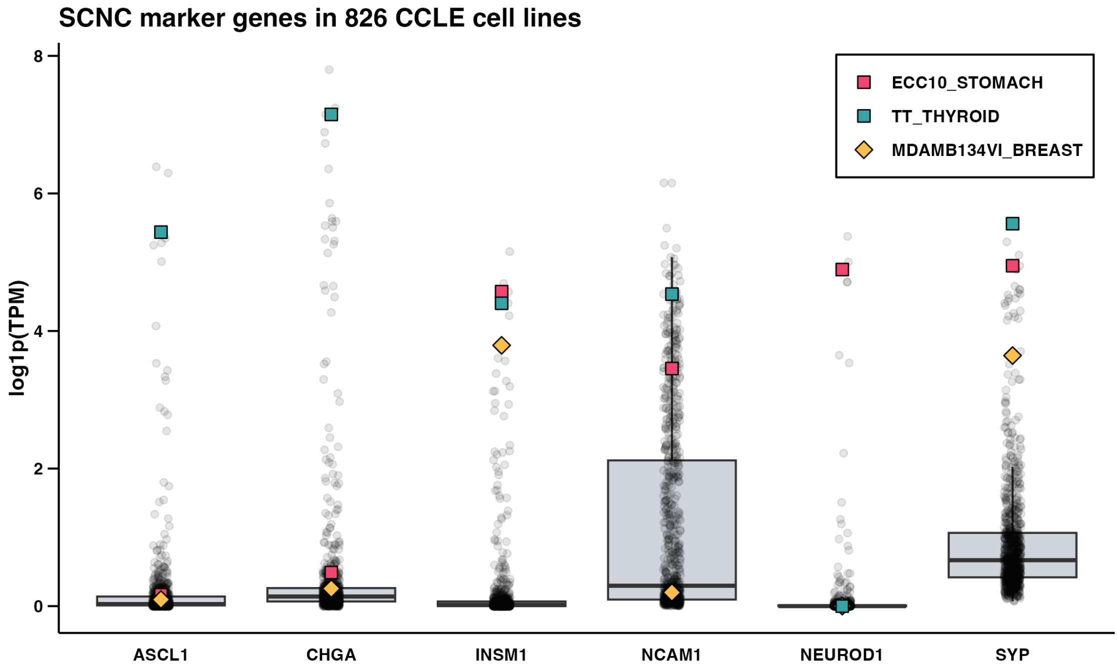
SCNC marker genes in 826 CCLE cell lines
log1p(TPM)
8
6
4
2
0
ECC10_STOMACH
TT_THYROID
MDAMB134VI_BREAST
ASCL1
CHGA
INSM1
NCAM1
NEUROD1
SYP

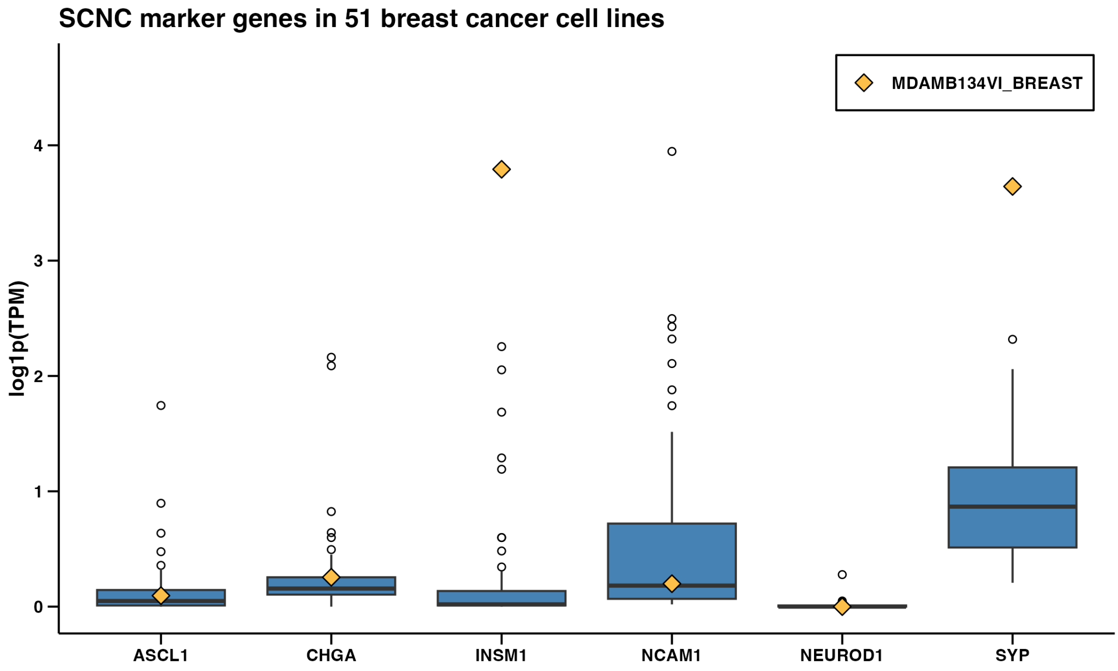
SCNC marker genes in 51 breast cancer cell lines
log1p(TPM)
4
3
2
1
0
MDAMB134VI_BREAST
ASCL1
CHGA
INSM1
NCAM1
NEUROD1
SYP

#### Supp. Fig 6

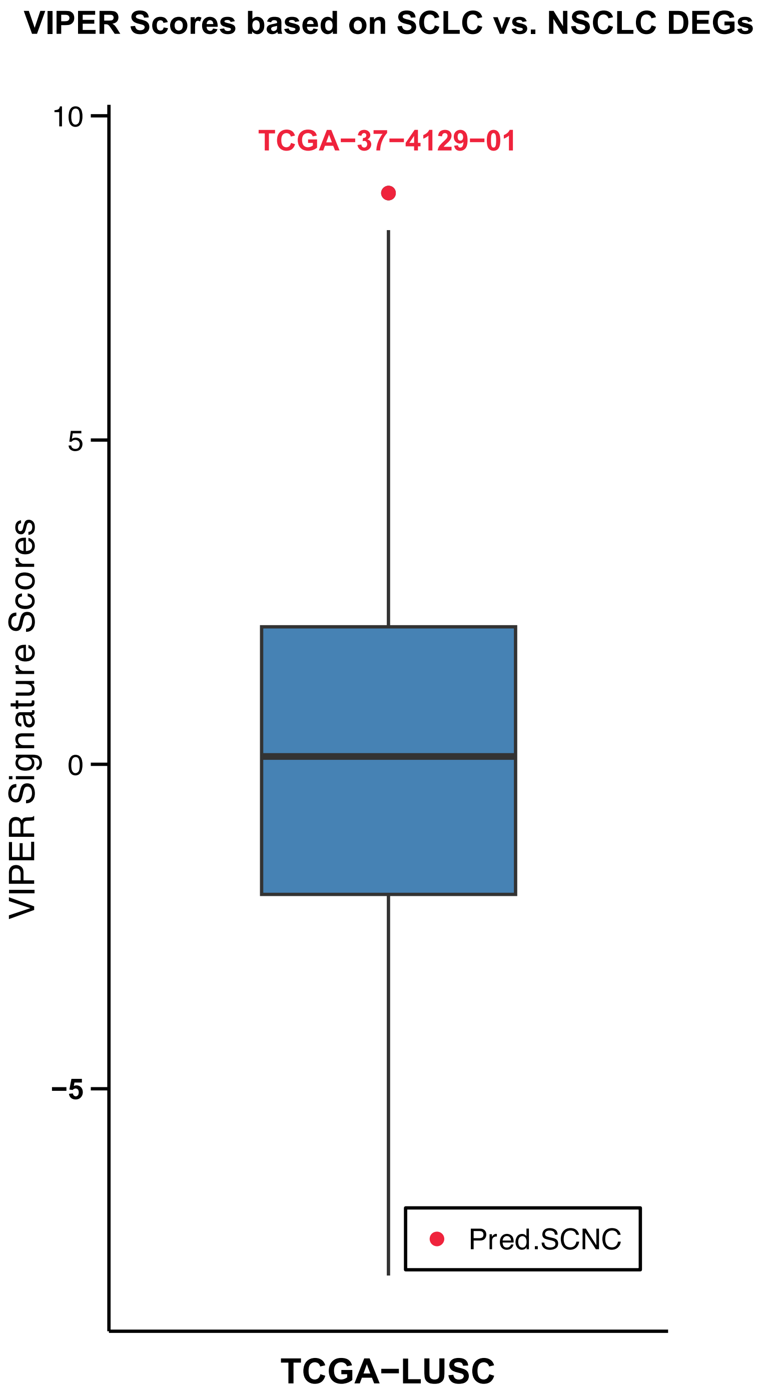

VIPER Scores based on SCLC vs. NSCLC DEGs
TCGA−37−4129−01
10
5
0
−5
VIPER Signature Scores
Pred.SCNC
TCGA−LUSC

**Supp. Fig 1.** APA Profiling in PDX model and health human tissues.488 lengthening and 122 shortening APA events were identified in the NEPC-PDX samples versus non-NEPC-PDX samples, under cutoffs: FDR<0.1, |FoldChange|>=1.4, diff_PDUI>=0.05 and min_PDUI>=0.15. 515 lengthening and 499 shortening APA events were identified in the healthy brain tissues versus other healthy tissues, under cutoffs: FDR<0.1, |FoldChange|>=1.4, diff_PDUI>=0.05 and min_PDUI>=0.15.

**Supp. Fig 2.** Upper panel: Heatmap showing the NA (missing value) ratio of 59 overlapped lengthening APA events across TCGA tumor types. Rows are grouped by NA ratio thresholds (NA ratio ⩾ 0.01 and < 0.01). The color scale represents NA frequency from 0 to 1. Lower panel: Box plots displaying NA frequency distribution for each of the 59 APA events across 25 TCGA tumor types (n = 7,659 samples). Each box plot represents the distribution of NA frequencies for one APA event across all tumor types. APA events are ordered by median NA frequency. The 20 events with consistently low NA frequencies (NA ratio < 0.01) were selected as conserved APA signatures for neural network model training.

**Supp. Fig 3.** Quality control of single-cell lung cancer. Single-cell RNA-seq data from 52 lung cancer samples were obtained from the Human Tumor Atlas Network (HTAN) database. Quality control filtering based on a minimal epithelial tumor cell abundance of 50 cells resulted in a high-quality cohort of 28 samples (22 SCNCs and 6 non-SCNCs) used for determining the optimal classification cutoff.

**Supp. Fig 4.** Copy number variation (CNV) analysis in SCNC and non-SCNC cells. CNV profiles across chromosomes in SCNC (SCLC) versus non-SCNC (NSCLC) cells from single-cell RNA-seq data. Heatmap shows inferred CNV scores for individual cells (rows) across genomic regions (columns). SCNC cells exhibit higher frequency and amplitude of copy number alterations compared to non-SCNC cells, indicating greater genomic instability that may contribute to the observed 3'UTR lengthening pattern and malignant phenotype.

**Supp. Fig 5.** Extended analysis of the expression level of SCNC marker genes within 826 cell lines and breast cancer cell lines separately. INSM1 and SYP rank the top in MDAMB134VI_BREAST among all breast cancer cell lines.

**Supp. Fig 6.** VIPER analysis validates predicted SCNC sample in TCGA-LUSC cohort. Distribution of VIPER scores in LUSC samples, with the predicted SCNC sample TCGA-37-4129-01 marked as an outlier, supporting its potential reclassification as an unannotated SCNC case within the TCGA-LUSC cohort.
